# Identification of a new mycobacterial peptide that controls the activity of the iron-dependent regulator, IdeR

**DOI:** 10.64898/2026.07.13.738276

**Authors:** Ashis Biswas, Assirbad Behura, Nishant Sharma, Shamba Gupta, Irene Pérez, Jesús Gonzalo-Asensio, Jun Yong Choi, G. Marcela Rodriguez

## Abstract

*Mycobacterium tuberculosis* (*Mtb*) must regulate intracellular iron to survive in the host and cause disease. IdeR (**I**ron-**de**pendent **R**egulator) is an essential *Mtb* protein that governs intracellular iron levels by regulating the expression of genes involved in iron metabolism. IdeR expression, however, is iron independent, and the mechanisms that regulate IdeR’s function are not fully understood. Here, we report the discovery of a previously unrecognized *Mtb* peptide (PRI) that regulates IdeR activity. PRI is induced under iron limitation; it binds to IdeR and restricts its activity. In addition, we demonstrate that altering the balance between PRI and IdeR impairs iron homeostasis and intracellular replication of *Mtb* in macrophages. The findings reveal a new paradigm in mycobacterial iron regulation and open new avenues for targeting iron homeostatic mechanisms in *Mtb*, which are crucial for virulence and antibiotic resistance.

## INTRODUCTION

Like most living organisms, *Mycobacterium tuberculosis* (*Mtb*), the bacterium responsible for human tuberculosis, utilizes iron as the preferred redox cofactor for proteins and enzymes essential for critical cellular functions, including energy metabolism and DNA replication. *Mtb* requires iron to replicate in the host and cause disease (1, 2). However, iron is not readily available in the host due to its insolubility under physiological conditions (3). To obtain iron, *Mtb* produces potent siderophores: mycobactin and carboxymycobactin (1, 4–8). These ferric iron chelators effectively compete with host proteins for iron (9). In addition, during macrophage infection, *Mtb* inhibits phagosome maturation while allowing phagosome fusion with early endosomes containing endocytosed transferrin, a reliable iron source (1, 5, 10, 11), and can also utilize iron in heme and hemoglobin (12–14).

Even though iron is vital, excess iron can be toxic because it readily catalyzes the production of harmful reactive oxygen species (ROS) from normal byproducts of aerobic respiration (15, 16) (17). Consequently, all aerobic organisms must tightly regulate intracellular iron to prevent oxidative stress. The Iron-dependent Regulator (IdeR) is a conserved protein in mycobacteria that governs iron homeostasis (18–23). IdeR functions as a metal and DNA-binding protein. Metal binding induces dimerization and conformational changes that activate DNA binding (24, 25). Two dimers of IdeR bind to each strand of a 19 bp palindromic DNA sequence (iron box) located upstream of iron-responsive genes, including those involved in iron sequestration, uptake, assimilation, and storage (18).

Mycobacterial genes encoding iron-acquisition-related proteins have a single iron box overlapping the promoter or the transcriptional start site (18). Consequently, IdeR binding to this sequence represses transcription. The genes encoding the iron storage proteins bacterioferritin (BfrA) and ferritin (BfrB) each contain tandem iron boxes upstream of their promoters (18, 21). IdeR binding to these sequences activates transcription, possibly by facilitating recruitment of RNA polymerase and, in the case of *bfrB,* by an anti-repressor mechanism that antagonizes the nucleoid-associated protein Lsr-2 (26).

IdeR is essential in *Mtb* (21). Deletion of *ideR* is lethal, and conditional depletion of IdeR results in deregulated siderophore synthesis, inadequate iron storage, accumulation of toxic intracellular iron, oxidative stress, and failure of *Mtb* to replicate in macrophages and *in vivo* in mice (22).

IdeR is well characterized structurally and functionally (21, 22, 25, 27–30). However, it remains unclear how its function is regulated. IdeR is constitutively expressed independent of iron conditions (21); the mechanisms that prevent IdeR from being activated by scarce iron in an iron-deficient bacterium and from prematurely repressing iron acquisition genes before the cell achieves iron sufficiency are not understood. This work identified a new *Mtb* peptide (PRI), upregulated during iron limitation, that interacts with IdeR and restricts its activity. We demonstrate that regulation of IdeR is crucial for adapting to iron limitation and that a balanced IdeR-PRI system is critical for *Mtb* replication in macrophages. The discovery of a mechanism of functional regulation of IdeR opens new avenues for developing antitubercular strategies that target iron homeostasis, thereby making *Mtb* more vulnerable to immune-mediated killing and antibiotics.

## RESULTS

### Identification of PRI

Detailed examination of the DNA sequence surrounding *ideR* identified an open reading frame encoding a putative 44-amino-acid-long peptide (based on the cut-off of 50 amino acids to be considered a protein) (Fig. 1A) corresponding to a small RNA previously sequenced in *Mtb* (31). This ORF is encoded in the opposite strand to *ideR,* overlapping the carboxy-terminal end of IdeR. Based on the results presented here, we have named the peptide encoded by this gene PRI, which stands for **P**eptide **R**egulator of **I**deR.

**Figure 1.**
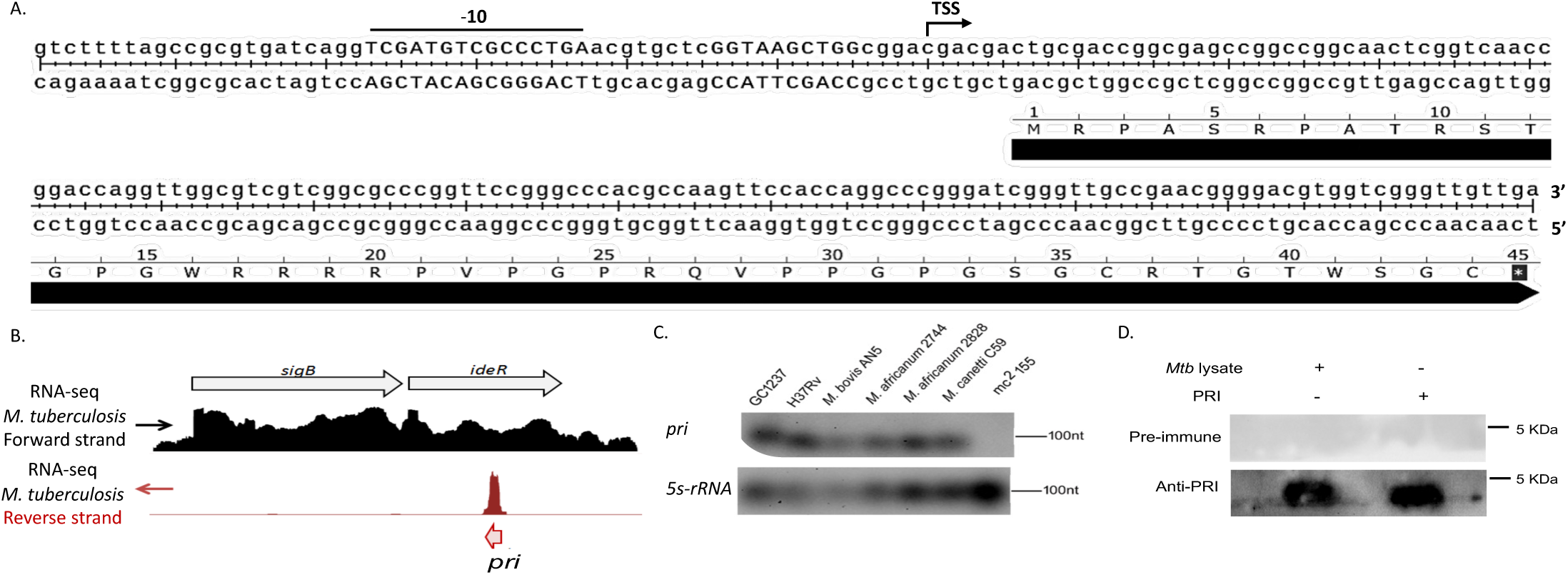
***Identification of PRI***: A. *pri* sequence; transcriptional start site (TSS) and the promoter sequence (-10) are indicated. B. RNAseq profile of *Mtb* showing *pri* transcript whose coordinates were found to coincide with those mapped experimentally. C. Northern Blot showing *pri* transcript and loading control 5S-rRNA in various mycobacteria. D. Western Blot of *Mtb* lysates with pre-immune control and immune anti PRI. Isolated PRI is included as a positive control.

The transcriptional start site of *pri* was identified by end joint mapping, as described in the Materials and Methods, and was found to coincide with the 5’ end determined by RNA sequencing. A 5’ untranslated region and a putative −10 sequence homologous to the consensus promoter sequence proposed by Cortes et al (32) were identified (Fig. 1A-B). Northern blot analysis confirmed a corresponding transcript in *Mtb* H37Rv, an *Mtb* clinical strain (GC1237), and other virulent mycobacteria, including *M. bovis*, *M. africanum,* and *M. canetti* (Fig. 1C). No transcript was detected in the non-virulent *M. smegmatis*.

PRI was visualized in *Mtb* cell extracts by Western Blot as a single band migrating identically to pure PRI at approximately 5 kDa, recognized by a specific antibody raised against chemically synthesized PRI but not by the pre-immune antiserum (Fig. 1D).

### PRI is induced in response to iron limitation

A possible connection between PRI and the *Mtb* response to iron was investigated by examining *pri* expression under a gradient of iron. *Mtb* was grown to the logarithmic phase in 7H10 agar and then transferred to iron-depleted minimal medium (MM) supplemented with increasing concentrations of iron. The abundance of the *pri* transcript was determined by qRT-PCR with specific primers (Table S2). The results showed an inverse correlation between iron in the medium and the abundance of the *pri* transcript (Fig 2). Induction of *pri* in low-iron conditions suggests that PRI might play a role in *Mtb*’s adaptive response to iron limitation.

**Figure 2.**
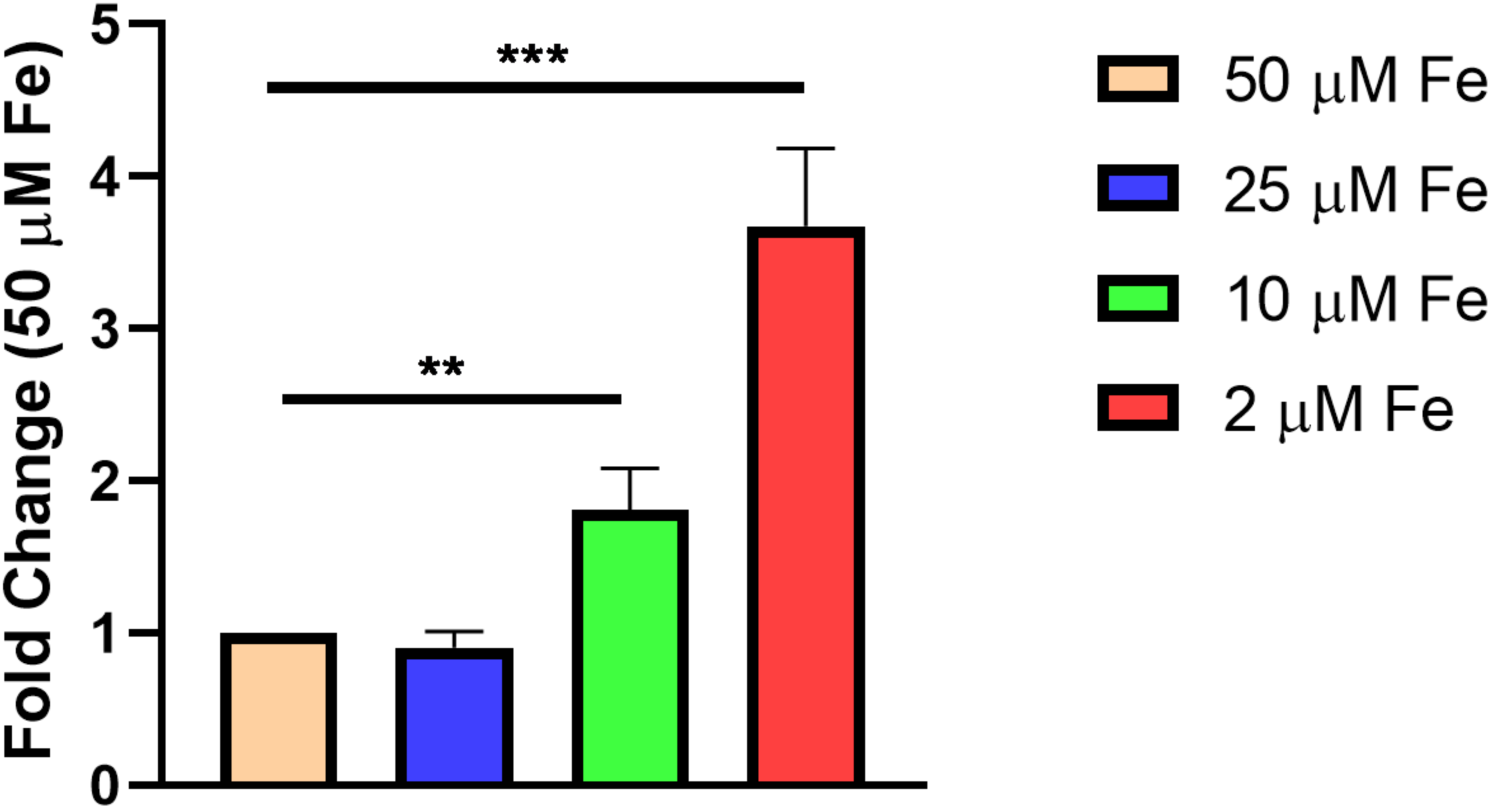
***Induction of pri by low iron.*** Expression of *pri* in *Mtb* H37Rv in iron-depleted medium repleted with increasing concentrations of FeCl_3_ was analyzed by qRT-PCR with Sybr green. The data are expressed as the fold change relative to 50 μM FeCl_3_ and presented as the mean ± SD from three biological replicates. \*\**p* < 0.01, \*\*\**p* < 0.001 Student’s *t*-test.

### Overexpression of PRI sensitizes *Mtb* to iron and oxidative stress

To investigate a possible role of PRI in the iron response, we generated Mtb-PRI^Tet,^ a strain of *Mtb* H37Rv transformed with a plasmid carrying a *pri* copy under a TetR-regulated promoter and induced in the presence of tetracycline (Fig S1A). To induce episomal *pri*, we used 1 μg.ml^-1^ of the tetracycline analog Anhydrotetracycline (Atc), which did not affect the growth of WT *Mtb* (Fig S1B) and assessed the effects of PRI overexpression on *Mtb* adaptation to diverse iron conditions. Atc addition inhibited the growth of Mtb-PRI^Tet^ in direct correlation with the concentration of iron in the medium (Fig 3A-C), indicating that augmented PRI sensitizes *Mtb* to iron toxicity. Mtb-PRI^Tet^ cultures treated with Atc also showed greater sensitivity to hydrogen peroxide than WT (Fig 3D). This is consistent with bacteria experiencing iron-fueled oxidative stress. Taken together, these results suggest that overexpressed PRI interferes with the safe handling of excess iron in *Mtb*.

**Figure 3.**
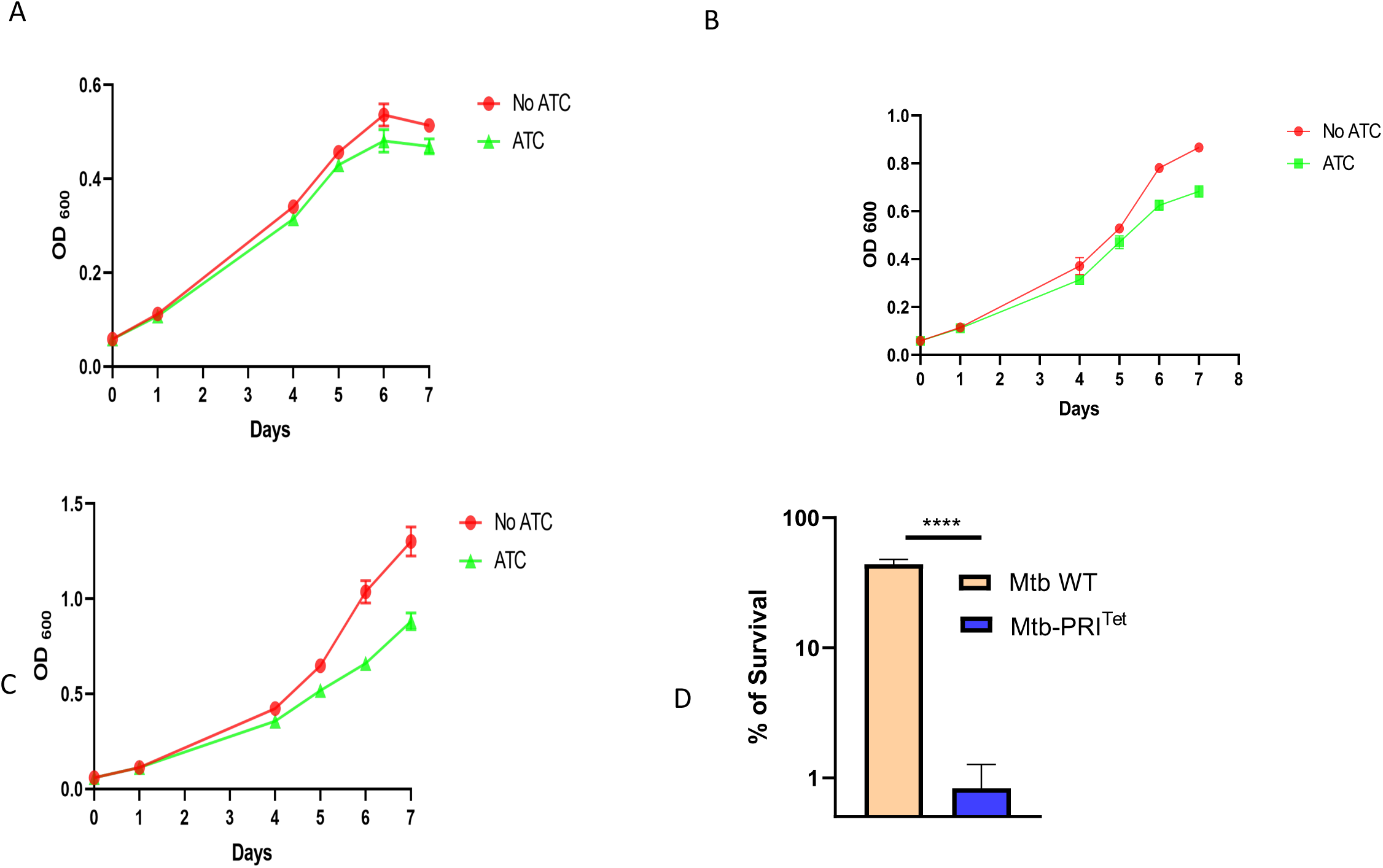
Effect of PRI overexpression on Mtb-PRI^Tet^ growth and oxidative stress sensitivity. A. Growth of Mtb-PRI^Tet^ in iron-depleted minimal medium supplemented with 2 μM FeCl3. B. Growth of Mtb-PRI^Tet^ in iron-depleted minimal medium supplemented with 10 μM FeCl_3_. C. Growth of Mtb-PRI^Tet^ in iron-depleted minimal medium supplemented with 50 μM FeCl3. D. Survival of WT and Mtb-PRI^Tet^ cultured in minimal medium supplemented with 50 μM FeCl_3_ and exposed to 1.25 mM H_2_O_2_ and Atc for 48 hours. The percentage of survival was determined from the number of CFUs recovered from cultures not exposed to H_2_O_2_ (100% survival). Data represent the mean ± SD from biological triplicates. **** *p*< 0.0001.

### PRI overexpression and regulation of IdeR-controlled genes

*Mtb* relies on IdeR for iron homeostasis and protection against iron-catalyzed oxidative stress (18–21). To determine whether overexpression of PRI leads to iron toxicity by interfering with IdeR, we compared IdeR transcript and protein levels in Mtb-PRI^Tet^ cultures with or without Atc. Even though no significant differences in IdeR levels were observed (Fig. 4), dysregulation of IdeR-regulated genes was apparent in cultures with Atc. The siderophore synthesis genes (*mbtB, mbtD, and mbtI)* and their positive regulator (*hupB*) (33), the iron-siderophore importer and ferric reductase-encoding gene *irtA* (2), the iron-siderophore utilization gene *eccA3* (34), and *ppe*37 were all derepressed, and *bfrB* (26) was poorly induced in high-iron medium (Fig 5). Atc did not affect the expression of IdeR-regulated genes in WT *Mtb,* ruling out that Atc signals changes in the expression of iron genes (Fig. S2). These results show that Atc-mediated overexpression of PRI negatively affects IdeR activity, both as a transcriptional repressor and as an activator.

**Figure 4.**
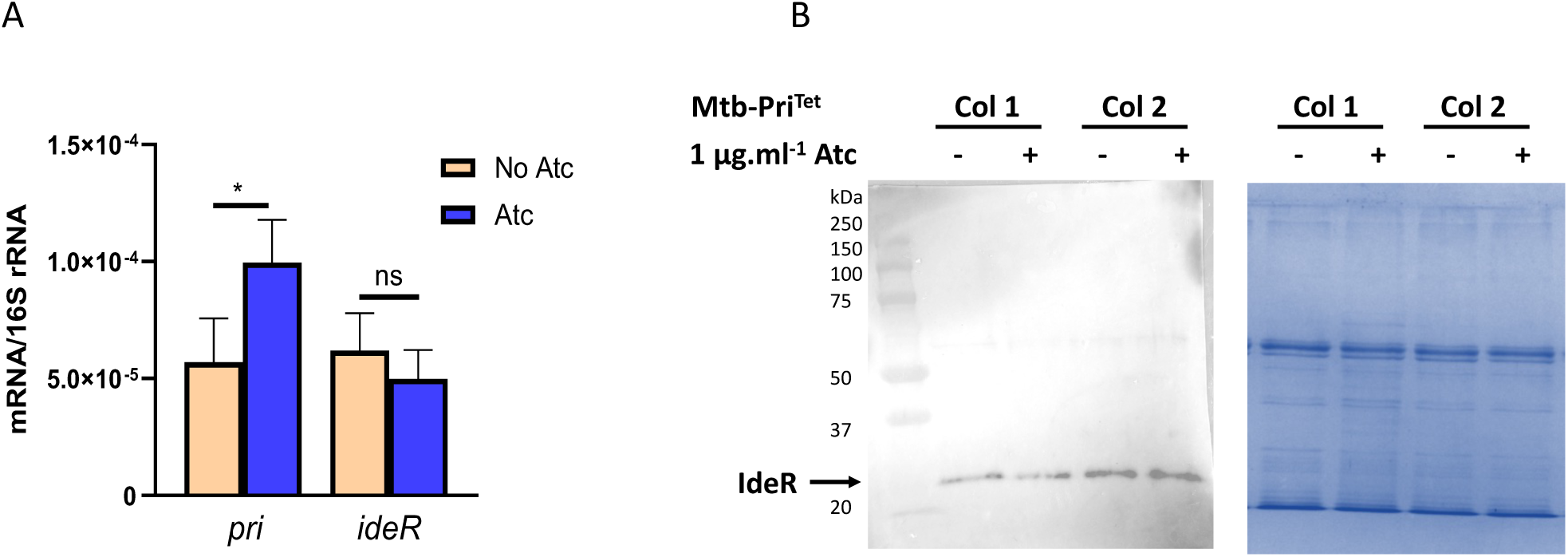
**IdeR expression in Mtb-PRI^Tet^**. A. Induction of *pri* in Mtb-PRI^Tet^ by Atc did not significantly alter *ideR* transcript. Transcripts were quantified by qRT-PCR with specific primers and normalized to 16s rRNA. Data are mean +/- SD.*p<0.05. B. The left panel shows a representative Western Blot (n=3) of two independent Mtb-Pri^Tet^ clones cultured ± Atc. IdeR was detected with a specific antibody (20). The right panel shows the corresponding Coomassie-stained gel confirming equal loading of bacterial lysates.

**Figure 5.**
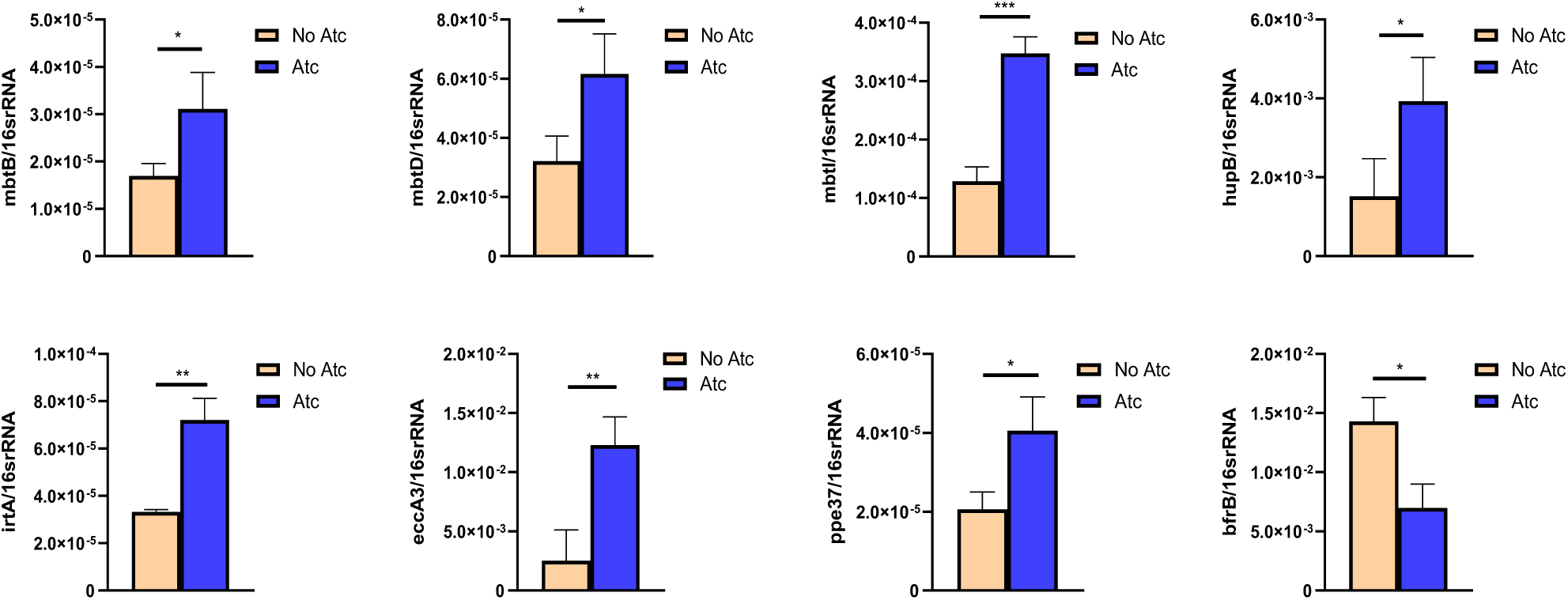
Expression of IdeR-regulated genes in cells overexpressing PRI. The graphs show the abundance of transcripts corresponding to genes normally repressed (*mbtB*, *mbtD*, *mbtI*, *hupB*, *irtA, ppe*37) and activated (*bfrB*) by IdeR under high-iron conditions. Mtb-PRI^Tet^ was cultured in high iron medium (50 μM FeCl_3_) with and without Atc, and gene transcripts were determined by qRT-PCR and normalized to 16S rRNA. Data are mean +/- SD.*p> 0.05; **p> 0.01; \*\*\**p*> 0.001 Student’s t-test. (n=3).

### PRI interacts with IdeR and inhibits its metal- and DNA-binding activities

To investigate the molecular mechanism by which PRI regulates IdeR activity, we tested a physical interaction between IdeR and PRI. Recombinant tag-less IdeR (Fig. S3) and PRI (Fig. S4) were purified, and their interaction *in vitro* was examined by Micro Scale Thermophoresis (MST). MST measures biomolecular interactions in solution by detecting changes in the movement of a target, in this case IdeR, along a temperature gradient as a function of a ligand concentration. The MST data indicate that PRI interacts with IdeR (K_d_ = 5.1± 0.89 μM), preferentially metal-free IdeR, as pre-incubation of IdeR with iron drastically reduced the interaction with PRI (Fig. 6). PRI-IdeR interaction in whole cells was confirmed by pull-down analysis; IdeR and PRI pulled down each other when co-expressed in *E. coli* (Fig 7).

**Figure 6.**
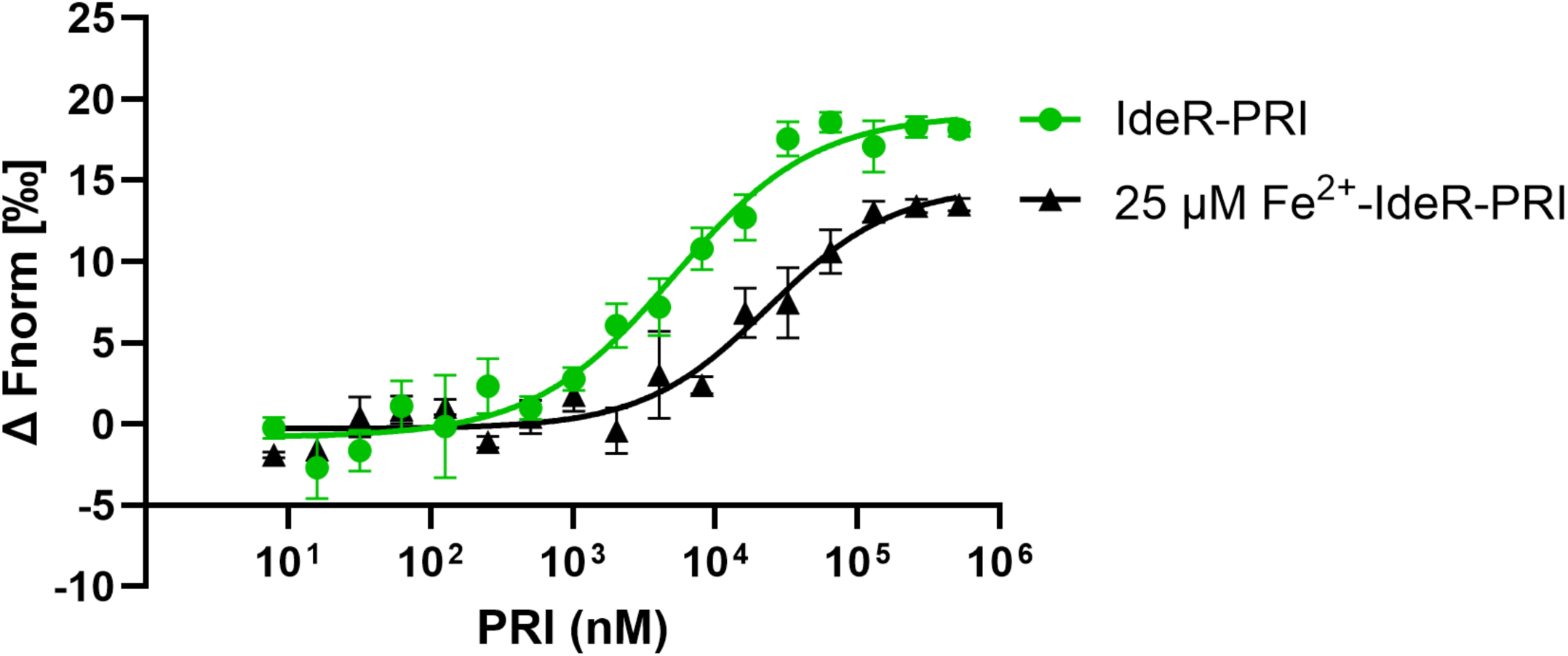
IdeR-PRI interaction measured by MST. MST signals of PRI titrated against 20 nM IdeR rendered metal-free by dialysis in Chelex-treated buffer containing EDTA. IdeR was incubated with 0 or 25 µM FeSO_4_ (as described in Materials and Methods) prior to measuring the interaction with PRI. A Kd=5.1±0.89 μM was calculated for the interaction between PRI and metal-free IdeR by Monolith Affinity Analysis Software.

**Figure 7.**
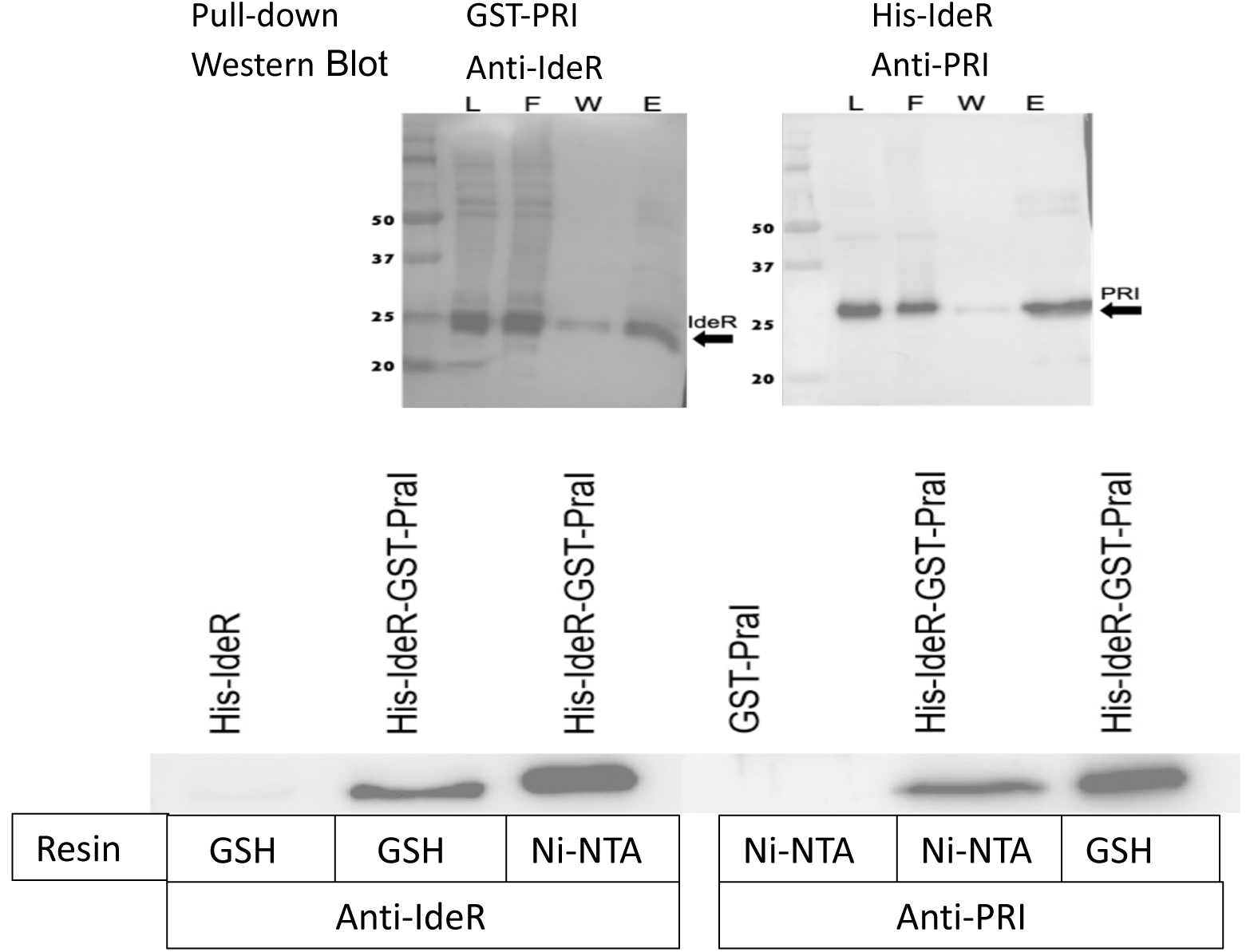
IdeR-PRI interaction in whole cells. Pull-down analysis of His-IdeR and GST-PRI co-expressed in *E. coli*. Left panel: The lysate was incubated with glutathione magnetic beads, and the load(L), flow-through (F), wash(W), and eluent(E) fractions were blotted with anti-IdeR antiserum. The arrow points to His-IdeR. Right panel: the lysate was incubated with Ni-NTA beads, and the L, F, W, and E fractions were blotted with Anti-PRI antiserum. The arrow points to GST-PRI. Lower panel: control that confirms no binding of purified His-IdeR to glutathione beads (GSH) or GST-PRI to Ni-NTA beads. The experiment was repeated three times.

Molecular dynamics (MD) simulation of PRI and molecular docking to IdeR suggested that PRI could interact with IdeR at the dimerization domain, which includes most of the metal-coordination residues. The N-terminal amine of M1 in PRI can form hydrogen bonds with IdeR’s H79 or C102, and R19 may form cation-ρε interactions with IdeR’s F128/H98 (Fig. 8). In addition, R6, R17, and R18 in PRI can form salt-bridge interactions with D17/E20, E95, and E96 of IdeR, respectively (Fig 7A-B). Lending support to this model of interaction, mutating R17, R18, and R19 in PRI to Alanine reduced PRI interaction with IdeR (Fig. 7c), and blocking or mutating IdeR’s unique Cys102, a key residue for metal-dependent activation of IdeR (29), abolished the interaction with PRI (Fig. 7D). These results suggest the interaction of PRI with IdeR may restrict metal binding and consequently DNA binding. In agreement with this notion, IdeR prebound to PRI exhibits reduced interaction with iron (Fig. 7E) and with DNA as detected by electrophoretic mobility shift assays (EMSA) (Fig 8A) and MST (Fig 8B) using as a probe a DNA fragment encompassing the iron boxes upstream of *bfrB*, labeled with Sybr Green or Cy5, respectively (26).

**Figure 8.**
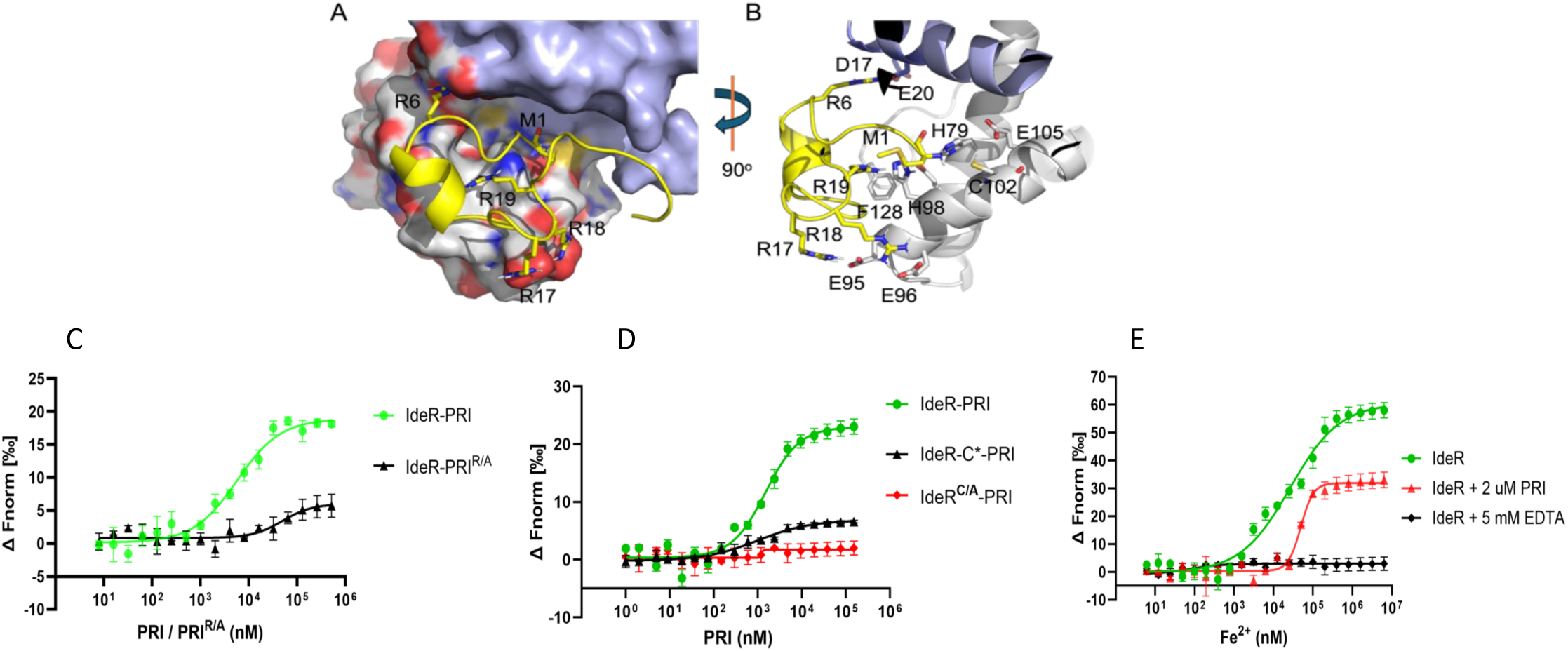
A-B. Model of PRI (yellow) interaction with IdeR at the secondary metal binding site obtained by MD. C. MST data of PRI or PRI with R17-20 mutated to Ala (PRI^R/A^) titrated against IdeR; D. MST data of PRI titrated against IdeR versus IdeR with C102 modified with maleimide (IdeR-C*) or mutated to Ala (IdeR^C/A^). E. Iron (FeSO_4_) titrated against IdeR or IdeR in complex with PRI in the absence or presence of EDTA as a control. Error bars represent the mean ± SD of technical triplicates. The experiments were repeated three times with independent preparations of purified proteins.

### Deregulated IdeR impairs *Mtb* growth under low-iron conditions

To assess the significance of regulating IdeR in *Mtb*, we engineered a system to overexpress IdeR without increasing PRI (since they share the same locus). We introduced at the *attB* site a chemically synthesized *ideR* copy in which the third position of all codons encompassing PRI was changed, leaving the amino acid sequence of IdeR intact while completely altering that of PRI. The resulting strain (ST448) has two functional copies of *ideR* and one (the native one) of *pri*. We also generated a control strain (ST459) that carries one extra copy of the intact *ideR/pri* locus at the *attB* site (Table 1). In both strains, the additional genes were expressed through their native promoters. Growth assays showed ST459 multiplied like WT *Mtb* independently of iron in the medium (Fig. 10B). In contrast, ST448 displayed a substantial growth defect in low-iron medium (Fig. 10A). This defect was partially alleviated by the addition of a high concentration of iron (150 μM), indicating that these cells are iron-deficient. These results show that tight regulation of IdeR is necessary for an effective adaptive response of *Mtb* to iron limitation.

**Figure 9.**
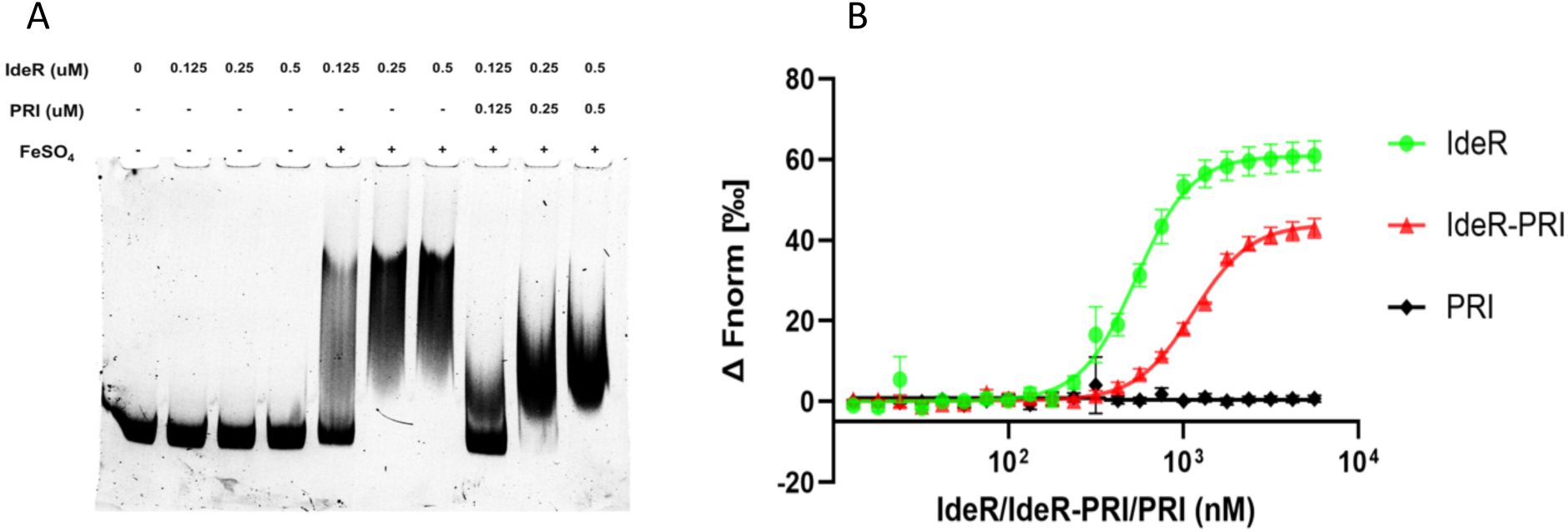
IdeR bound to PRI shows reduced DNA binding activity. A. Electrophoretic mobility shift assay (EMSA) of a DNA fragment encompassing the IdeR binding site upstream *bfrB* incubated with IdeR with or without FeSO_4_ and with or without PRI. Protein-DNA complexes were resolved by electrophoresis, and the DNA was visualized by staining with Sybr Green. The experiment was repeated three times with independent isolates of IdeR and PRI. B. MST data measuring the interaction of IdeR versus IdeR-PRI and PRI as control with the same DNA probe used in EMSA labeled with Cy5 during synthesis. Error bars represent the mean ± SD of technical triplicates. The experiments were repeated three times with independent protein preparations.

**Figure 10.**
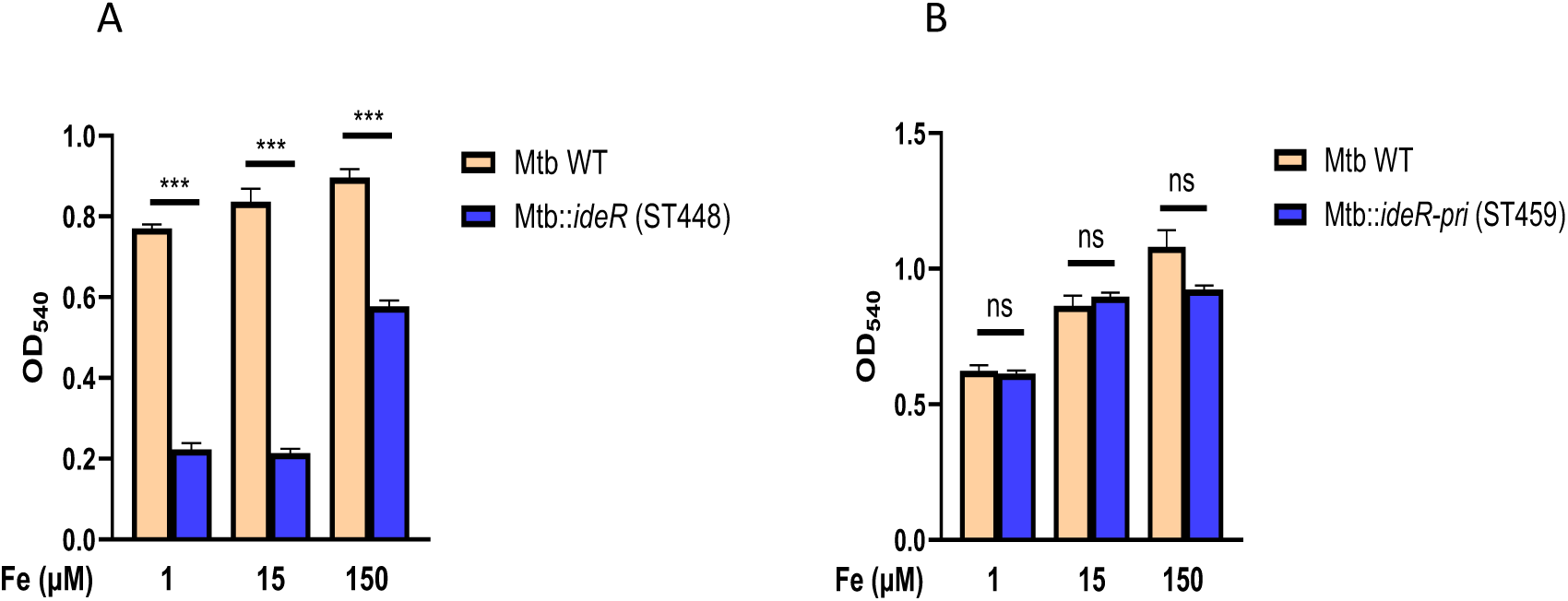
Overexpression of IdeR impairs *Mtb* growth under low-iron conditions. WT and *ideR* diploid *Mtb* strain (ST448) in (A) and *ideR-pri* diploid strain (ST459) in (B) were inoculated in an iron-defined medium at an initial O.D_540_ of 0.05 and incubated at 37 °C. The graph shows the O.D 540 nm on day 4. Data are mean +/- SD.* *p $ 0.01,* \*\**p $ 0.001,* *** *p ≤ 000.1* (n=3).

**Figure 11.**
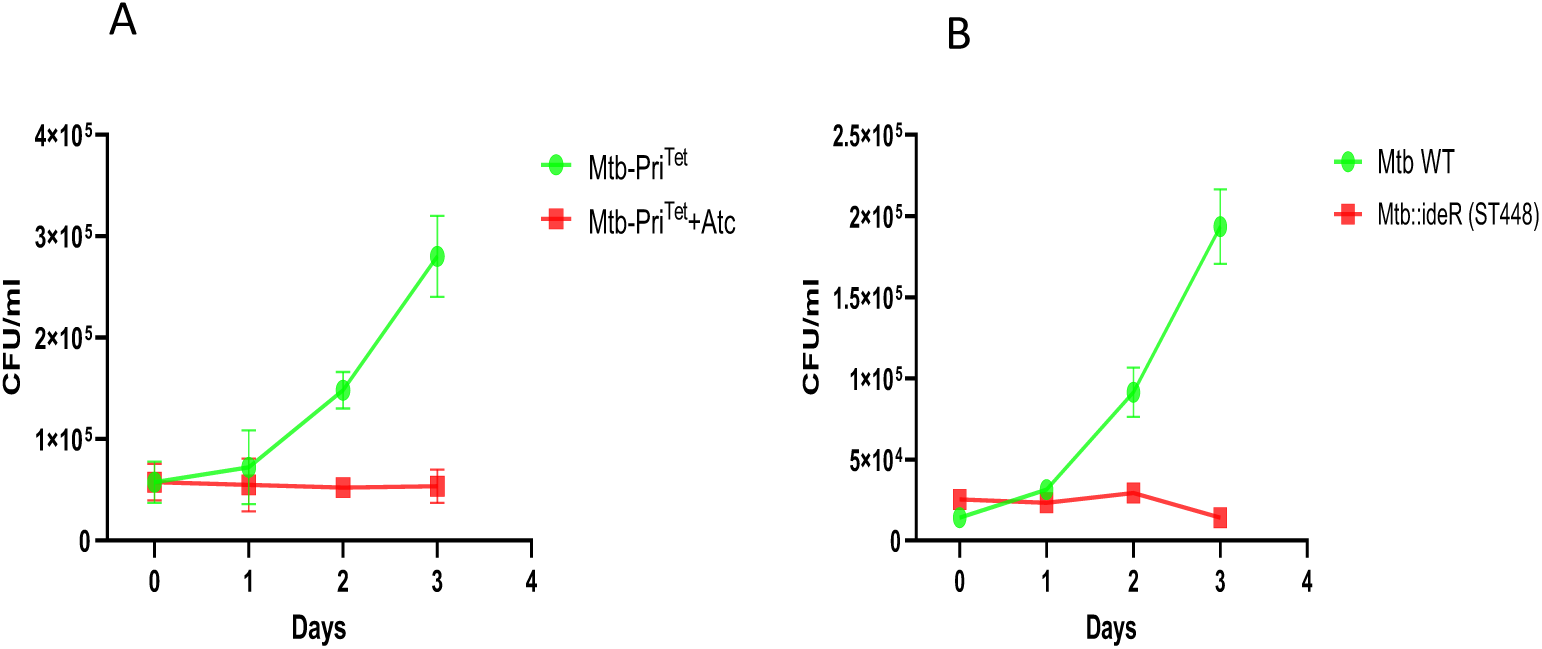
Intracellular Growth of *Mtb* overexpressing PRI or IdeR. A. Growth of Mtb-PRI^Tet^ in THP-1 cells ± Atc. B. Growth of *Mtb* WT and the strain having an additional copy of *ideR* in the chromosome (ST448) in THP-1 cells. Error bars represent the mean ± SD of technical triplicates. The experiment was repeated twice.

#### An unbalanced PRI-IdeR system attenuates Mtb growth in macrophages

We next determined the impact of altering the normal balance between IdeR and PRI on *Mtb* adaptation to the phagosome environment by assessing replication of IdeR or PRI-overexpressing bacteria in macrophages. THP-1 cells differentiated into macrophages were infected with WT, ST448, or Mtb-PRI^Tet^in the presence or absence of Atc, and bacterial replication was evaluated by enumerating CFUs at the indicated times. Although infection efficiency was equivalent across strains (time 0), intracellular replication of ST448 and Mtb-PRI^Tet^ in the presence of Atc was drastically reduced compared to WT *Mtb*. A control experiment showed that Atc had no effect on the replication of WT *Mtb* in THP-1 cells (Fig. S6). These results indicate that altering the IdeR-PRI system—either by increasing IdeR, as in ST448, or by enhancing PRI, as in Mtb-PRI^Tet^ significantly hinders *Mtb*’s ability to adapt to the intracellular environment and proliferate in macrophages.

## DISCUSSION

Although IdeR has been extensively studied, the mechanisms underlying its regulation have remained unclear. IdeR expression is not regulated by iron (21), which raises the question of what prevents IdeR from binding scarce iron and prematurely repressing iron-acquisition genes before the cell satisfies its iron needs. The current paradigm assumes that IdeR binds only surplus iron. However, given that IdeR can compete with iron metalloenzymes for iron, and that IdeR activation in an iron-deficient bacterium could lead to premature shutdown of iron acquisition and compromise bacterial growth, it is likely that a mechanism exists to ensure IdeR activation occurs only when the cell has achieved iron sufficiency. In agreement with the need to regulate IdeR, we observed that doubling the *ideR* gene dose was sufficient to hamper *Mtb* adaptation to iron limitation and the phagosome environment.

This study identified PRI, a previously unrecognized peptide upregulated under low-iron conditions, that can regulate IdeR function. Overexpression of PRI recapitulated hallmark phenotypes of IdeR depletion, including iron-metabolism gene dysregulation accompanied by iron sensitivity, and oxidative stress (22). Regulation of IdeR by PRI was supported by the observed physical interaction between these two proteins *in vitro* and in whole cells.

IdeR has a high-affinity metal binding site (MBS1) and an auxiliary site (MBS2). Iron binding at MBS1 is sufficient to stabilize IdeR dimers, but metal binding to MBS2 is necessary for activating DNA binding (35). Our MST analysis shows that PRI interacts preferentially with metal-free IdeR; however, they do not rule out an interaction with IdeR in which the high-affinity MBS1 is occupied. Nonetheless, our computational modeling and mutagenesis analysis suggest that PRI interaction with IdeR requires C102, a critical metal-ligand residue at MBS2. Based on our results, we propose a model in which PRI restricts metal binding at MBS2, effectively elevating the threshold of iron required to fully activate IdeR for DNA binding. In agreement with this model, MST results show reduced IdeR-metal interaction when IdeR is bound by PRI compared to free IdeR, and a significant inhibition of IdeR-DNA complex formation in the presence of PRI.

Current studies are focused on resolving the structure of the IdeR in complex with PRI, which will clarify the molecular mechanism by which PRI regulates IdeR activity and generate a framework for rational design of inhibitors of IdeR or the PRI-IdeR interaction.

Consistent with our model of IdeR regulation by PRI, increasing IdeR without a commensurate increase in PRI impeded an adequate adaptive response to iron limitation, reflected in deficient growth under those conditions. It is likely that PRI regulation of IdeR not only facilitates robust expression of the iron acquisition machinery during iron deficiency but also prevents upregulation of ferritin in low-iron conditions, which would be detrimental during iron limitation, as ferritin could compete with iron-requiring enzymes for scarce iron.

The results of this study highlighted the importance of a balanced IdeR-PRI system for intracellular replication of *Mtb*, as overexpression of either IdeR or PRI inhibited *Mtb* growth in THP-1 cells. This is consistent with previous studies demonstrating that both siderophore-mediated iron acquisition(2) and proper iron storage in ferritin are necessary for *Mtb* to replicate in naive macrophages (36).

It has not been possible to inactivate *pri*, even in an *ideR* diploid strain, suggesting PRI may be essential in *Mtb.* Current studies are focused on generating a conditional PRI mutant to dissect the effects of PRI depletion during infection.

Previously, we demonstrated that ferritin is crucial for *Mtb* to tolerate antibiotics (36). PRI offers a new platform to design IdeR inhibitors that, by interfering with IdeR-mediated activation of *bfrB,* enhance *Mtb* susceptibility to existing antibiotics.

In sum, the discovery of PRI deepens our understanding of the mechanisms underlying iron homeostasis in *Mtb* and is expected to inspire research on novel therapeutic interventions that target essential iron-regulatory pathways to enhance immune- and antibiotic-mediated killing of *Mtb*.

## MATERIAL AND METHODS

### Bacterial strains, media, and growth conditions

Bacterial strains and plasmids used in this study are listed in Table 1. *Mtb* H37Rv (ATCC) was used in all experiments. *Escherichia coli* was routinely grown at 37 °C in Luria-Bertani (LB) medium. *Mtb* strains were recovered from frozen stocks on Middlebrook 7H10 agar (Difco) supplemented with 0.5% v/v glycerol, 0.05% tween 80, and 10% v/v albumin-dextrose-NaCl complex (ADN). Fe-depleted minimal medium (MM) was used for the growth of *Mtb* under iron-defined conditions. MM contains 0.5% (w/vol) asparagine, 0.5% (w/vol) KH_2_P0_4_, 2% glycerol, 0.05% Tween-80, and 10% ADN; pH: 6.8. MM was treated with 5% Chelex-100 (BioRad) for 24 hours, at 4 °C to remove traces of iron. Chelex-100 was removed by filtration through a 0.22 μm filter, and the medium was supplemented with 0.5 mg. L^-1^ of ZnCl_2_, 0.1 mg.L^-1^ of MnS0_4_, and 40 mg.L^-1^ of MgS0_4_. The amount of residual Fe in this medium, determined by atomic absorption spectroscopy, is less than 1 µM.

When required, antibiotics were added at a final concentration of 100 g.ml^-1^ of carbenicillin (Car), 50 or 20 g.ml^-1^ of Kanamycin (Kan) for *E. coli* and mycobacterial cultures, respectively, 75 μg.ml^-1^ of Spectinomycin (Spec), and 20 μg.ml^-1^ of Streptomycin (Strp) and, unless specified, 1μg.ml^-1^ Anhydrotetracycline.

### Plasmid construction

A tetracycline-inducible expression plasmid was constructed by inserting the Tet repressor sequence from pST-KT (37) in the SnaBI site of a pST-2K (37) derived plasmid in which the *Sph*I site at position 763 was mutated by overlapping PCR, generating pSM947. To generate pSM1095, the *pri* gene was PCR-amplified from *Mtb* chromosomal DNA using specific primers (Table S1) and cloned into the SphI-NotI sites of pSM947. pSM1100 was produced by site-directed mutagenesis of pSM1095. To generate the *ideR* merodiplid strain (ST448), we constructed pSM1053 by cloning a synthetic IdeR gene into the StuI site of pSM1049, which is derived from pSM316 after removal of the kanamycin resistance cassette. In the synthetic *ideR*, the third position of all codons in *pri* has been changed to preserve the amino acid sequence of IdeR intact while mutating all residues in PRI.

pSM305 was generated by PCR amplification of *ideR* and *pri* with their respective promoters, followed by cloning the product into pMV306.

To express recombinant IdeR, *ideR* was PCR amplified from the Mtb and cloned at the NdeI-HindIII sites of pET28TEV (38) to generate pSM918, which carries IdeR tagged at the amino-terminal with a 6xHis tag cleavable by the TEV protease.

To express recombinant PRI, the *Mtb pri* sequence was PCR-amplified and cloned into the BamHI site of pQlinkG to generate pSM1096, which carries an in-frame gene fusion of PRI with an amino-terminal GST tag cleavable by the TEV protease.

### RNA extraction and Northern blot analyses

Mycobacterial cultures were grown to the late exponential phase (OD_600_=0.5-0.6) and pelleted by centrifugation. Bacteria were resuspended in 1 ml RNA Protect Bacteria Reagent (Qiagen), incubated for 5 min at room temperature, and then centrifuged. Bacterial pellets were resuspended in 0.4 ml lysis buffer (0.5% SDS, 20 mM Na Acetate, 0.1 mM EDTA) and 1 ml phenol: chloroform 1:1 (pH=4.5). Suspensions were transferred to tubes containing glass beads (Qbiogene) and lysed using a RiboLyser (Fastprep instrument) with a three-cycle program (15 sec at speed 6.5 m), with cooling on ice for 5 min between pulses. Samples were then centrifuged, and the homogenate was transferred to a tube containing chloroform: isoamyl alcohol 24:1. Tubes were inverted carefully before centrifugation, and the aqueous phase was then transferred to a fresh tube containing Na-acetate (0.3 M), pH=5.5, and isopropanol. Precipitated nucleic acids were collected by centrifugation. The pellets were rinsed with 70% ethanol and air-dried before resuspension in RNase-free water. DNA was removed by 1 h treatment with DNase (Ambion) at 37°C. RNA integrity was evaluated by agarose gel electrophoresis, and the absence of contaminating DNA was confirmed by the lack of amplification products after 30 PCR cycles.

Northern blot was performed using the DIG Northern starter kit (Roche) following the manufacturer’s recommendations. Briefly, total RNA, 100 ng for 5S rRNA and 10 µg for mRNA detection, were separated by electrophoresis in a 2% agarose, 2% formaldehyde gel in 1X MOPS buffer. RNA was transferred by capillary blotting onto Hybond-N+ nylon membranes (Amersham), UV-crosslinked, and incubated with the desired probe. Digoxigenin (DIG)-labeled probes were synthesized by in vitro transcription, following the manufacturer’s directions to detect *rrf* (5S rRNA) and PraI transcripts using the primer pairs NB-5S-rRNA-fw and NB-T7-5S-rRNA-rv, NB-T7-as-ideR-Fw/NB-as-*ideR*-rv or NB-T7-Ms-as-ideR-Fw/NB-Ms-as-ideR-rv and ncRv2711c-PBF and ncRv2711c-PBR for *M. smegmatis* and *M. tuberculosis* complex (MTBC) strains, respectively (Table S1). RNA transcripts complementary to each probe were detected by Western blot using an anti-DIG antibody conjugated to alkaline phosphatase and the chemiluminescent substrate CDP-*Star*. The size of the transcripts was estimated by comparison with the DynaMarkerTM Pre-stain Marker for Small RNA Plus.

### Transcriptional start site point mapping

To identify the 5’ and 3’ ends of PraI transcript, we adapted a procedure that uses T4 RNA ligase to circularize RNAs followed by cDNA synthesis, PCR amplification, and DNA sequence of joint ends (39): total RNA was ligated by T4 RNA ligase 1 (New England BioLabs) to produce circular RNA. 5 μg RNA was ligated in a total volume of 20 µl including 2 µl 1X T4 RNA ligase buffer, 2 µl RNase OUT (ThermoFisher Scientific), 2 µl RNA T4 ligase, 0.1 µl ATP, and DEPC treated water to 20 µl. The mixture was incubated for 2 h at 25 °C, and the reaction was stopped by incubation at 65 °C for 10 min. Circular RNA was purified by chloroform extraction and ethanol precipitation. The RNA pellet was dried and re-suspended in 5 µL DEPC water. cDNA synthesis was conducted using 1 μl of a 10 μM solution of a gene-specific primer, mixed with 5 μg RNA and 2 μl dNTP mix, and the volume was adjusted to 12 μl with DEPC-treated water. RNA and primer were denatured at 65°C for 5 min and then placed on ice for 5 min. 8 μl of a master reaction mix including 4 μl 5X cDNA synthesis buffer, 1μl DTT (0.1M), 1 μl RNaseOUT^TM^, 1 μl DEPC-treated water, and 1 μl ThermoScript^TM^ RT enzyme was added to the denatured ligated RNA and incubated at 60°C for 60 min. The reaction was terminated by incubation at 85°C for 5 min. Finally, RNA was degraded with RNase H (1 μl) for 20 min at 37°C. The cDNA containing the 5’-3’ junction of *pri* was amplified with internal specific primers (PRIc-1 and PRIc-2) in a 50 μl reaction volume in a Mastercycler gradient thermocycler (Eppendorf) at the following conditions: denaturation at 94°C for 5 min, followed by 35 cycles of 94°C for 1min, 58°C for 45 sec, and 72°C for 1 min and ended with a final extension for 10 min at 72°C. After PCR amplification, the products were resolved on a 2% agarose gel, and a ∼100 bp band was excised and purified using the MinElute® Gel Extraction Kit (Qiagen) according to the manufacturer’s instructions. The purified PCR product was cloned into the PCR-Blunt II-TOPO Vector (Thermo Fisher Scientific) according to the manufacturer’s instructions, and transformants were selected with Kanamycin (Kan). Recombinant plasmids from 3 Kan-resistant transformants were extracted and sequenced. The determined sequences were used as queries on the BLAST server of the National Center for Biotechnology Information (NCBI) using BLASTN to identify the junction point between the two opposite ends of the RNA. This method was validated with known bacterial sRNA. As experimental control in parallel to the ends of *pri,* we mapped the ends of Mcr7, a well-characterized *M. tuberculosis* sRNA (40).

### Western Blot

*Mtb* was grown in the indicated conditions to the logarithmic phase. Protein extracts were generated by harvesting the bacteria by centrifugation, washing the pellet twice with PBS, resuspending it in 1ml Tri-Reagent (MRC), and preparing cell lysates by bead-beating in a FastPrep-24 instrument (MP Biomedicals, Santa Ana, CA) with 200 μm zirconia-silica beads (Benchmark Scientific). The aqueous phase containing RNA was separated from the lysates using 100 ml of BCP (Molecular Research Center), followed by centrifugation at 12,000 xg, at 4°C for 10 min. The organic phase was removed, and 300 μl of ethanol was added to precipitate the DNA. DNA was removed by centrifugation at 2,000 xg at 4 °C for 5 min. The supernatant was transferred to a fresh tube, and proteins were precipitated with 1.5 ml isopropanol by centrifugation at 12,000 xg at 4 °C for 5 min.

Protein pellets were washed twice by adding 2 mL 0.3 M guanidine hydrochloride solution, incubating at RT for 20 min, and pelleting down at 12,000 ×g at 4°C for 5 min. The pellets were washed by adding 2 ml of ethanol, incubating at room temperature for 20 min, then centrifuging at 12,000 g, 4°C for 5 min. The supernatant was discarded, and pellets were air-dried and resuspended in 1% SDS. Proteins were resolved by SDS-PAGE, transferred to a nitrocellulose membrane in an iBLOT3, and detected with our anti-IdeR (20) diluted 1:1000 or anti-PRI (1:5000). Anti-PRI was produced in rabbits immunized with chemically synthesized PRI (Thermo Fisher). Anti-rabbit IgG-HRP was used as a secondary antibody, and detection was performed using enhanced chemiluminescence (ECL) (GE Healthcare).

### Quantitative RT-PCR

cDNA was synthesized from RNA (1 µg) with SuperScript-IV Reverse Transcriptase (Thermo Fisher Scientific) and random hexamer primers. The reaction was incubated for 10 min at 23°C, 12 min at 52°C, and 10 min at 80°C. The cDNAs were processed for real-time PCR using TB Green *Premix Ex Taq* II (Takara) (30 s at 95°C; 40 cycles of 5 s at 95°C, 30 s at 60°C) on an Agilent AriaMx real-time PCR System (Agilent) using specific primers (Table S1). All samples were normalized against the transcript levels of 16S ribosomal RNA. Relative mRNA levels were determined by the comparative CT method (41).

### Sensitivity to Hydrogen Peroxide

*Mtb* strains were cultured in MM with 50 μM FeCl_3_ to logarithmic phase, diluted to an OD_540_ of 0.05, and incubated in the same medium with or without 1.25 mM H_2_O_2,_ in the presence or absence of Atc (1 μg.ml^-1^) for 48 hr. At the end of this time period, an aliquot of each culture was diluted and plated to determine the number of survivors as CFUs.

### Protein purification

*IdeR purification*: pSM918 was transformed into the *E. coli* strain C41(DE3). A Kan^R^ *E. coli* transformant was grown overnight in 2 ml of LB containing 50 μg/mL of kanamycin (LB-Kan) at 37°C and 200 rpm agitation. This culture was used to inoculate 1 L of LB-Kan and grown at 37°C and 200 rpm to an OD_600_ of 0.8. Cells were then incubated overnight at 16°C and 200 rpm with 400 µM IPTG. Induced cultures were centrifuged at 6000 x g at 4°C, and the cell pellet was resuspended in 20 ml of 50 mM Tris-HCl with 150 mM NaCl (lysis buffer) supplemented with 20 μl of 2 mg.ml^-1^ DNase (Sigma) and a cocktail of protease inhibitors (Roche). Cells were lysed in a French Press (1500 psi) for 3 cycles. The cell lysate was centrifuged at 20,000 x g for 45 min, at 4°C, and the supernatant was collected and loaded onto a nickel column (His-Trap) (Thermo Fisher) pre-equilibrated with lysis buffer. Unbound material was removed by washing the column with 50 ml of lysis buffer, followed by a 30-100 mM Imidazole gradient in 50 mM Tris-HCl and 500 mM NaCl. IdeR was eluted from the column with 10 ml of 250 mM Imidazole in 50 mM Tris-HCl and 500 mM NaCl.

Fractions containing His-IdeR (determined by SDS-PAGE) were pooled and dialyzed against 50 mM Tris-HCl, 50 mM NaCl with 1 mM DTT at 4 °C overnight. The His tag was removed by digestion with 800 μg TEV protease overnight at 4 °C. The His-tag-free IdeR was separated from non-cleaved His-IdeR using a second His-Trap column, which was washed with 10 mL of 50 mM Tris-HCl, 500 mM NaCl. The flow-through and first wash were collected and concentrated in a 10 KDa Amicon concentrator. IdeR was then dialyzed against 50 mM Tris-HCl, 50 mM NaCl, 1 mM DTT, and 1 mM EDTA at 4 °C overnight and further purified by FPLC in a Superdex 200 Increase 10/300 GL gel filtration column using as running buffer 50 mM Tris-HCl, 50 mM NaCl,1 mM EDTA and 1 mM DTT. All solutions used for protein purification were treated with Chelex-100 and sterile-filtered before use. The protein was concentrated in a 10 KDa Amicon concentrator, aliquoted, and stored at −80 °C (Supplementary Fig). The Cys102Ala mutant of IdeR was generated by Quick Change mutagenesis (Stratagene), cloned into pET28TEV, expressed in *E. coli,* and purified as described for wild-type IdeR.

*PRI purification:* PRI expression plasmid pSM1096 was transformed into the *E. coli* strain C41(DE3). A Kan^R^ *E. coli* transformant was grown for 4 h in 100 ml of LB containing 100 μg.ml^-1^ of carbenicillin (LB-Car) at 37°C and 200 rpm agitation. This culture was split into 2 flasks, each one containing 2 L of LB-Car, and grown for 3-4 hours at 37°C with continuous shaking at 200 rpm. The cells were grown to an O.D_600_ of 0.8 and then induced with 400 µM IPTG overnight at 16 °C and 200 rpm. Induced cultures were centrifuged at 6000 x g at 4°C, and the cell pellet was resuspended in 50 ml of 50 mM Tris-HCl, pH 8.0, with 500 mM NaCl (lysis buffer) supplemented with 20 μl of 2 mg.ml^-1^ DNase and a cocktail of protease inhibitors (Roche). Cells were lysed in a French Press (1500 psi) for 3 cycles, and the cell lysate was centrifuged at 20000 x g for 1 h to remove cell debris. The supernatant was then incubated with 1 ml of Glutathione Sepharose 4B beads (Cytiva) in lysis buffer overnight at 4°C in a roller mixer. The beads were collected in a Poly-Prep chromatography column (Bio-Rad) and washed with 200 ml lysis buffer. GST-PRI was eluted with 30 mM reduced-glutathione prepared in lysis buffer, with the pH adjusted to 9.0 with 10 M NaOH. The eluted GST-PRI was then dialyzed against 1 L of 20 mM Tris-HCl, 50 mM NaCl, 1 mM DTT, and 1 mM EDTA overnight. The dialyzed protein was concentrated to 3-5 ml in a 10KDa Amicon Ultra. The concentrated protein was digested with 800 μg of TEV protease at 4°C overnight. The tag-free PRI was collected in a Centriprep-30 (Amicon) by vacuum suction, aliquoted, and stored at −70°C.

### Electrophoretic Mobility Shift Assays (EMSA)

A DNA fragment encompassing 500 bp upstream of *bfrB* was PCR amplified, purified by agarose gel extraction, and used as a probe. IdeR-DNA binding reactions included 30 ng of the DNA probe, IdeR, and binding buffer [20 mM Tris-HCl (pH 8.0), 1 mM DTT, 50 mM KCl, 5 mM MgCl_2_, 0.05 mg ml^-1^ BSA, and 10% glycerol] in a final volume of 20 μl. The reaction was initiated by adding 0.2 mM FeSO_4_ and allowed to proceed at room temperature for 30 minutes. At the end of the incubation, the sample was mixed with 6X loading buffer containing 10 mM Tris-HCl (pH 7.6), 0.03% bromophenol blue, and 60% v/v glycerol, and loaded onto a 6% polyacrylamide gel containing 40 mM Tris-acetate (pH 8.0). The gel was run at 110V at 4 °C, then stained with Syber Green (1X in Tris-acetate buffer) (Invitrogen) and visualized using a GelDoc imager (BioRad).

To assess the effect of PRI, the DNA probe was incubated with equimolar concentrations of IdeR and PRI in the binding buffer for 10 min before adding FeSO_4_. The reaction, electrophoresis, and visualization of the protein-DNA complex were carried out as described above.

### Microscale Thermophoresis

*IdeR labeling:* IdeR was labeled with RED NHS 2nd Generation or RED Maleimide 2nd Generation (Nanotemper) according to the manufacturer’s instructions. Prior to labeling, the IdeR solution was buffer-exchanged using the supplied desalting column, and then 100 μL of 10 μM IdeR protein was mixed with 5 μL reconstituted dye (25 μL DMSO per vial). The labeling reaction was allowed to proceed for 30-45 minutes in the dark at RT, and the product was purified using the size-exclusion column provided in the kit. Protein concentration and degree of labeling (DOL) were measured using NanoDrop spectroscopy at 205 nm and 650 nm, respectively. Calculations were done using NanoTemper DOL calculator. Typically, the recovered concentration was 100-200 nM and the DOL 0.66-1.12. Protein solution was aliquoted and kept at –70 °C, and the working concentration of labeled protein was 20 nM.

*Reaction preparation:* To obtain a binding isotherm 16 sequential dilutions of the tested ligand were made and mixed with 20 nM IdeR and binding buffer (20 mM Tris-HCl,12.5 mM KCl, 5 mM MgCl_2_,10% Glycerol, 1 mM DTT, 50 μg/mL BSA, pH 8.0 with or without iron as indicated) in a final vol of and 25 μl. All reactions were carried out in low-binding tubes to prevent sample loss. The reaction components were mixed well by pulse centrifugation, and then incubated at room temperature for 30 min. Reactions were then loaded into a premium capillary (Nanotemper) and read on a Monolith NT.115 (Nanotemper), which measures the fluorescence change upon the introduction of infrared radiation for 20 seconds. Readings were performed at 22°C with a medium MST and an excitation power in a range from 20% to 50%. The data were analyzed using MO Affinity Analysis (x86) software on the Monolith NT.115.

When measuring DNA binding by MST, a 5’ Cy5-labeled dsDNA encompassing the IdeR binding region upstream of *bfrB* (Integrated DNA Technologies) was incubated at a final concentration of 25 nM with IdeR, PRI, or a mix of the two using the same binding buffer as in EMSA. The RED filter option of Monolith NT.115 from Nano Temper Technologies was used for this assay. Data analysis was performed by MO Affinity Analysis (x86) software.

A computational modeling pipeline was implemented after MST data collection to obtain full titration curves for downstream nonlinear regression analysis. Simulations and curve fitting were done using Python 3.11 software, using SciPy and NumPy for the nonlinear regression, computations, and replicate variation. The Fnorm experimental results for each titration curve were then analyzed using a 4PL fit, which yields the sigmoid shape expected from the thermophoresis assay. The model is defined as:

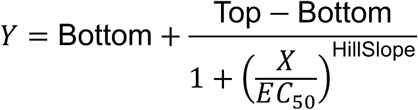

The optimization algorithm for nonlinear least squares was implemented via curve fitting with scipy.optimize.curve fit in Python, enabling unconstrained estimation of the parameters Bottom, Top, EC₅₀, and HillSlope from experimental data. Experimentally determined MST values were used to estimate these parameters. To simulate realistic variation observed during MST experiments, the model-derived MST values were replicated three times by adding normally distributed noise (5%) to the predicted MST signals using numpy.random.normal. Thus, the randomness present in MST experimental data, as well as the overall shape of the sigmoid curve, were captured. Replicate simulations were performed only for concentrations where experimental MST values were missing, resulting in uniform replicate counts across all titration concentrations. The merged dataset thus obtained was then subjected to nonlinear regression analysis.

### Pull-down analyses

E. coli C41 was transformed with pSM918 and pSM1096. Double transformants were selected in Kan plus Car, and one clone was grown to an OD_600_ of 0.6 and induced with 0.4 mM of IPTG for 4 hours. Cells were collected by centrifugation, suspended in PBS containing a protease inhibitor cocktail (Roche), and lysed in a bead beater for 2 cycles of 1 min each. Cell lysates were cleared from cell debris by 30 min centrifugation at 13000×g at 4°C. A portion of the lysate derived from *E. coli* expressing His-IdeR was added to HisPur-NiNTA agarose beads (ThermoScientific) and another portion to Glutathione Sepharose 4B (Cytiva) to control for nonspecific binding. Similarly, a portion of the lysate derived from *E. coli* expressing GST-PRI was added to Glutathione Sepharose 4B, and another portion was added to HisPur-NiNTA agarose beads to control for nonspecific binding. Lysates derived from *E. coli* expressing both His-IdeR and GST-PRI were also added separately to HisPur-NiNTA and Glutathione Sepharose 4B resins and incubated overnight at 4°C with gentle agitation. Beads and bound material were recovered by centrifugation and washed twice with PBS with 0.1% Triton-X100. Material bound to HisPur-NiNTA was eluted with 50 mM Tris-HCl, 500 mM NaCl, 300 mM Imidazole, pH 8.0, while that bound to Gluthatione Sepharose 4B was eluted with PBS with 30 mM reduced Glutathione, pH 9.0. Aliquots of the load, wash, and eluted material were loaded into a 15% SDS-PAGE. The gel was transferred to a PVDF membrane, processed for Western Blotting, and stained with anti-IdeR (1:1000) or anti-PRI (1:5000). Anti-rabbit IgG-HRP-linked was used as the secondary antibody, and detection was performed using enhanced chemiluminescence (ECL) (GE Healthcare).

### Macrophage infection

THP-1 cells were infected as previously described (2). Briefly, 1×10 ^^5^ THP-1 cells per well were induced to differentiate into macrophages with 50 nM phorbol myristate acetate for 24 hr. The differentiation medium was replaced with medium containing Mtb at a multiplicity of infection of 1 bacterium per 50 macrophages. After 3 hours of incubation at 37 °C in a 5% CO_2_ atmosphere, the medium was removed, and the cells were washed extensively with pre-warmed PBS to remove extracellular bacteria. Finally, 100 μl of warm RPMI with or without Atc (1 μg.ml^-1^) was added to each well. The plate was incubated at 37°C in a 5% CO_2_ atmosphere. RPMI with or without Atc was replaced every 48 hrs. At indicated time points after infection, macrophages were lysed with 0.05% sodium dodecyl sulfate (SDS). Serial dilutions of the lysates were made and plated in 7H10. CFU were counted after 4 weeks of incubation at 37°C.

### Statistical Analysis

All experiments were performed in triplicate. Data are expressed as mean ± SD. Differences between frequencies were assessed by the Student’s t-test (bilateral and unpaired) using a P value of <0.05 as statistically significant.

## Acknowledgments

We thank Long Xiang Xie and Hamel Amin for their assistance in this project.

