## Supplemental material for "Identification of a new mycobacterial peptide that controls the activity of the iron-dependent regulator, IdeR"

SUPPLEMENTAL INFORMATION

| | Characteristics | Reference or source |
|---|---|---|
| **Plasmids** | | |
| pMV306 | Integrative mycobacterial vector | {Stover, 1993 #1467} |
| pSM316 | Integrative mycobacterial vector | {Manganelli, 2004 #1712} |
| pSM305 | *ideR* & *pri* with their own promoters cloned into PMV306 | This work |
| pSM918 | pET28TEV carrying His-IdeR. | {Kurthkoti, 2015 #2101} |
| pSM947 | pST-2K with the Tet repressor from pST-KT inserted at the SnaBI site and the Sph I site at position 763 removed. | {Parikh, 2013 #1999} and this work |
| pSM1053 | *ideR* sequence with missense mutations in *pri* cloned in pSM316 modified to remove the Kan resistance cassette. | This work |
| pSM1095 | *pri* cloned in pSM947 | This work |
| pSM1096 | pQlinkG expressing GST-PRI | This work |
| pSM1100 | *pri* with R17-20A mutations cloned in pSM947 | This work |
| pSM1104 | *ideR with C102A mutation in pSM918* | This work |
| **E. coli strains** | | |
| Top 10 | F⁻ mcrA Δ(mrr-hsdRMS-mcrBC) ϕ80lacZΔM15 ΔlacX74 recA1 araD139 Δ(ara-leu)7697 galU galK rpsL(StrR) endA1 nupG. | Thermo Fisher |
| C41(DE3) | | Sigma Aldrich |
| **Mtb Strains** | | |
| H37RV | Wild-type Mtb | ATCC |
| ST448 | H37Rv transformed with pSM1053 | This work |
| ST453 | H37Rv transformed with pSM1095 | This work |
| ST459 | H37Rv transformed with pSM305 | This work |

**Supplementary Table 1.** Plasmids and bacterial strains used and generated in this work.

| Primers | Sequence 5' to 3' | Purpose |
|---|---|---|
| PRIc-1 | CCTGGTCCGGTTGACCGA | TSS mapping |
| PRIc-2 | TTGGCGTCGTCGGCGCCCGGTT | TSS mapping |
| *bfrB*Rev | GCAGAACCACTGCATGAAC | qRT-PCR reverse primer for *bfrB* |
| *bfrB*Fw | GGAAACCAGTTCGACAGACC | qRT-PCR forward primer for *bfrB* |
| *mbtB*Rev | TAATAAGGTCAACGCAAGTTCG | qRT-PCR reverse primer for *mbtB* |
| *mbtB*qRTFw | GCGACTTTCCCATCAGTGTT | qRT-PCR forward primer for *mbtB* |
| *irtA* Rev | GAGTCGCCGATTAGCAGATAC | qRT Reverse primer for *irtA* |
| *irtA* Fw | AAACCTGGCGCAACCATA | qRT forward primer for *irtA* |
| *mbtD* Rev | CAGGTTTGCGTACCAGTAGTC | qRT Reverse primer for *mbtD* |
| *mbtD* Fw | GCTGCCTGACTCCGAATTTA | qRT forward primer for *mbtD* |
| *mbtI* Rev | CGATGTCGGTAATCTCCTCAAG | qRT Reverse primer for *mbtI* |
| *mbtI* Fw | CTCGTGATGACCTGGAATCAA | qRT forward primer for *mbtI* |
| *eccA3* Rev | AATAGCGGCTCTACCGATCTA | qRT Reverse primer for *eccA3* |
| *eccA3* Fw | GCTGGACCATTCCGAACAT | qRT forward primer for *eccA3* |
| *ppe37* Rev | GCGAACGGTCCGAAATAGAT | qRT Reverse primer for *ppe37* |
| *ppe37* Fw | CAGGTGTTGGACTGGTTCAT | qRT forward primer for *ppe37* |
| *hupB* Rev | CTTAACAGCGGGTCCTTCTG | qRT Reverse primer for *hupB* |
| *hupB* Fw | GCGAGACAGTAAAGGTGAAGC | qRT forward primer for *hupB* |
| *ideR* Rev | GGTTTCGACGGTTACTCGTG | qRT Reverse primer for *ideR* |
| *ideR* Fw | GGTCAAGGTGCTCAACAACC | qRT forward primer for *ideR* |
| *pri* Rev | GGGCCTGGTGGAACTTG | qRT Reverse primer for *pri* |
| *pri* Fw | CCGGCAACTCGGTCAAC | qRT forward primer for *pri* |
| *16s rRNA* Rev | CGGCTGCTGGCACGTAGTTG | qRT Reverse primer for *16s rRNA* |
| *16s rRNA* Fw | ATGACGGCCTTCGGGTTGTAA | qRT forward primer for *16s rRNA* |

**Supplementary Table 2.** Primer sequences used in this work.

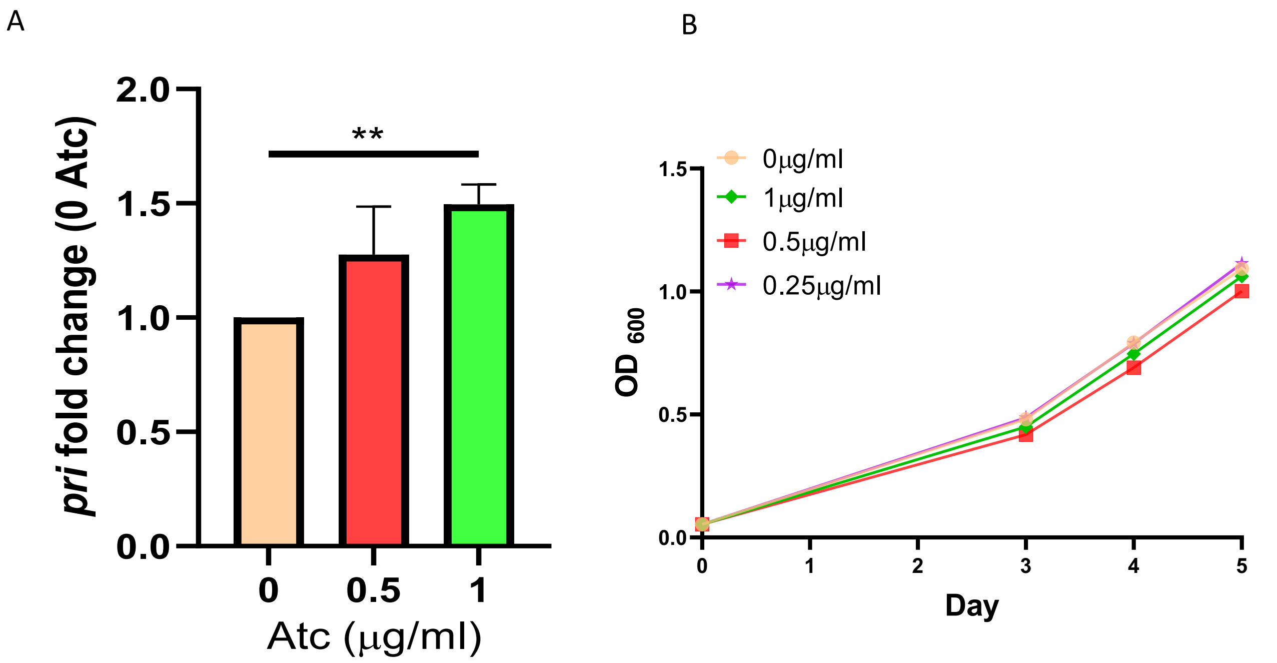


**Supplementary Figure 1.** A. Induction of *pri* in Mtb-PRI$^{Tet}$ by Atc. The *pri* transcript was detected by qRT-PCR with Syber green. The data are expressed as the fold change relative to 0 µg Atc and presented as the mean ± SD from three biological replicates. $^{**}p < 0.01$ B. Growth of WT *Mtb* H37Rv in medium containing the indicated concentrations of Atc. The experiment was repeated twice with technical triplicates.

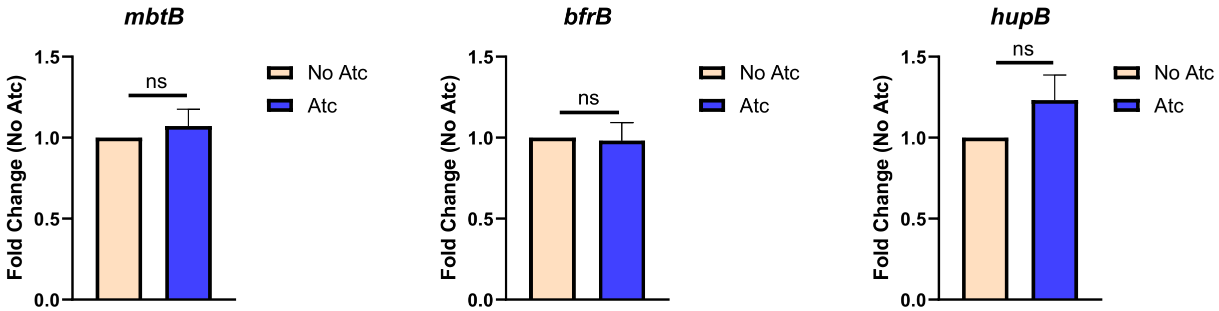


**Supplementary Figure 2. Expression of IdeR-regulated genes in WT *Mtb* treated with Atc.** Atc (1 $\mu g.ml^{-1}$) had no significant effect (NS) on the expression of the IdeR-regulated genes in WT *Mtb*. Abundance of gene transcripts was determined by qRT-PCR. The data are expressed as the fold change relative to no Atc and presented as the mean ± SD from three biological replicates.

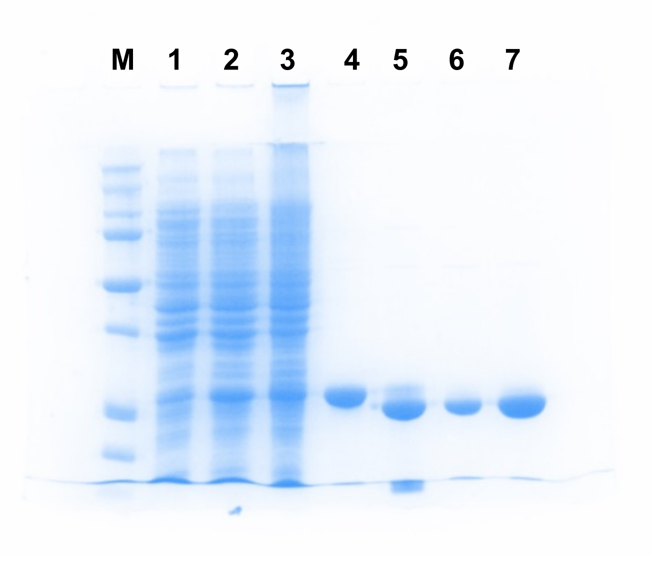


**Supplementary Figure 3. Purification of recombinant IdeR.** His-tagged IdeR was overexpressed and purified from *E. coli* by Nickel affinity chromatography. M-Molecular weight markers, 1- Uninduced *E. coli* lysate; 2- IPTG induced *E. coli* lysate; 3-Unbound flow through; 4- Imidazole eluted His-IdeR; 5- TEV cleaved His-IdeR; 6-7 Tag-free purified IdeR.

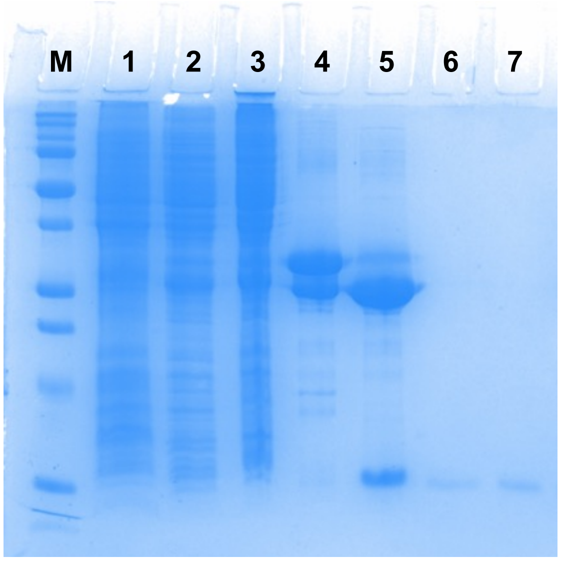


**Supplementary Figure 4. Purification of recombinant PRI.** Glutathione-S transferase (GST) -tagged PRI was overexpressed and purified from *E. coli* by affinity chromatography. M-Molecular weight markers, 1- Uninduced *E. coli* lysate; 2- IPTG induced *E. coli* lysate; 3-Unbound flow through; 4- glutathione eluted GST-PRI; 5- GST TAG cleavage; 6-7 Tag-free purified PRI.

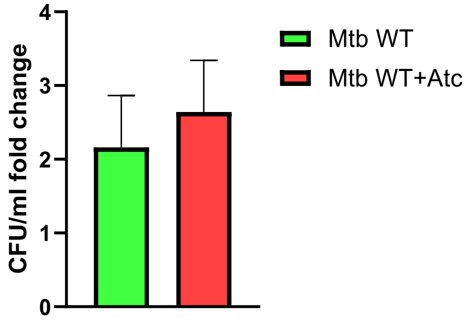


**Supplementary Figure 5. Atc does not affect the replication of WT *Mtb* in THP-1 cells.** THP-1 cells infected with WT Mtb were treated or not with Atc 1 μg. ml-1, and bacterial load was determined 2 hr (time 0) and 3 days post-infection by lysing the macrophages, plating dilutions of the lysate, and counting CFUs. The data represent the fold change of bacterial number at day 3 post-infection.
